# Fermented food metagenomics reveals substrate-associated differences in taxonomy, health-associated- and antibiotic resistance-determinants

**DOI:** 10.1101/2020.03.13.991653

**Authors:** John Leech, Raul Cabrera-Rubio, Aaron M Walsh, Guerrino Macori, Calum J Walsh, Wiley Barton, Laura Finnegan, Fiona Crispie, Orla O’Sullivan, Marcus J Claesson, Paul D Cotter

**Affiliations:** Teagasc Food Research Centre, Moorepark, Fermoy, Cork, Ireland; APC Microbiome Institute, University College Cork, Cork, Ireland; School of Microbiology, University College Cork, Cork, Ireland

## Abstract

Fermented foods have been the focus of ever greater interest as a consequence of purported health benefits. Indeed, it has been suggested that the consumption of these foods that help to address the negative consequences of ‘industrialization’ of the human gut microbiota in Western society. However, as the mechanisms via which the microbes in fermented foods improve health are not understood, it is necessary to develop an understanding of the composition and functionality of the fermented food microbiota to better harness desirable traits. Here we considerably expand the understanding of fermented food microbiomes by employing shotgun metagenomic sequencing to provide a comprehensive insight into the microbial composition, diversity and functional potential (including antimicrobial resistance, carbohydrate-degrading and health-associated gene content) of a diverse range of 58 fermented foods from artisanal producers from around the Globe. Food type, i.e., dairy-, sugar- or brine-type fermented foods, was to be the primary driver of microbial composition, with dairy foods found to have the lowest microbial diversity. From the combined dataset, 127 high quality metagenome-assembled genomes (MAGs), including 10 MAGs representing putatively novel species of *Acetobacter, Acidisphaera, Gluconobacter, Lactobacillus, Leuconostoc* and *Rouxiella*, were generated. Potential health promoting attributes were more common in fermented foods than non-fermented equivalents, with waterkefirs, sauerkrauts and kvasses containing the greatest numbers of potentially health-associated gene clusters (PHAGCs). Ultimately, this study provides the most comprehensive insight into the microbiomes of fermented foods to date, and yields novel information regarding their relative health-promoting potential.

**Importance:** Fermented foods are regaining popularity in Western society due in part to an appreciation of the potential for fermented food microbiota to positively impact on health. Many previous studies have studied fermented microbiota using classical culture-based microbiological methods, older molecular techniques or, where deeper analyses have been performed, have involved a relatively small number of one specific food type. Here, we have used a state-of-the-art shotgun metagenomic approach to investigate 58 different fermented foods of different type and origin. Through this analysis, we were able to identify the differences in the microbiota across these foods, the factors that drove their microbial composition, and the relative potential functional benefits of these microbes. The information provided here will provide significant opportunities for the further optimisation of fermented food production and the harnessing of their health promoting potential.

## Introduction

Fermentation is a form of food preservation with origins that can be traced back to the Neolithic age[1]. Despite recent advances in food preservation and processing, fermentation continues to be widely used as a means of preservation and is the focus of renewed interest due to increased appreciation of the organoleptic, nutritive and, especially, health promoting properties attributed to many fermented foods[2, 3].

Indeed, various fermented foods have been shown to have enhanced attributes relative to the corresponding raw ingredients by virtue of the microbial metabolites produced[4–8], the removal of allergens[9], other desirable biological activities[10, 11] and/or containing microbes that have the potential to confer benefits following consumption[12, 13]. Furthermore, although antibiotic use, sanitation and food processing have greatly reduced the number of deaths due to infectious diseases, these activities have also minimised our exposure to microbes and are thought to have contributed to the ‘industrialisation’ of the human microbiome and associated increases in chronic diseases[14, 15]. It has been suggested that fermented foods offer a means of safe microbial exposure to compensate for the absence/removal of desirable host microbes[15, 16].

Due to these potential benefits, and an increasing appreciation that the study of these foods provide valuable fundamental insights into simple microbial communities[17, 18], developing an even greater understanding of the microbiology of these foods has the potential to be of considerable value.

Advances in high throughput sequencing technology have revolutionised the study of microbial populations, including those present in foods. Although, to date, the vast majority of studies relating to fermented foods have employed amplicon sequencing to study bacterial and fungal composition[19–36], there have been some exceptional studies in which shotgun sequencing has been employed to gain a greater insight into the taxonomy and functional potential of specific fermented foods[37–49]. Despite this, studies across a broad variety of such foods using this approach have been lacking to date. Here we address this issue by employing shotgun metagenomic sequencing to investigate the microbiota of broad range of, including many previously unexplored, fermented foods.

## Results

### Fermented food microbiomes can be distinguished on the basis of substrate type

Shotgun metagenomic sequencing of 58 food samples (347,841,507 total reads; with an average of 5,997,267 reads per sample) and associated metadata (i.e., country of origin [‘country’], specific source of product [‘producer’], presence/absence of starter culture [‘fermentation’], solid or liquid foods [‘state’] and [’substrate’]) (**Table 1**), revealed that the microbiomes of these foods most significantly clustered on the basis of food substrate (i.e., dairy, such as kefir and cheese; brined, such as sauerkraut and kimchi; sugar, such as kombucha and water kefir; **Table 2**, **Figure 1**). Ten characteristics of the food microbiome were defined and differences across these characteristics were statistically examined (**Table 2**); 4 taxonomic levels (species, genus, family and phylum), 4 functional profiles (Superfocus 1,2 and 3, and Carbohydrate functions, which were a subset of Humann2 output), the bacteriocin profile and the antimicrobial resistance profile.

**Figure 1.**
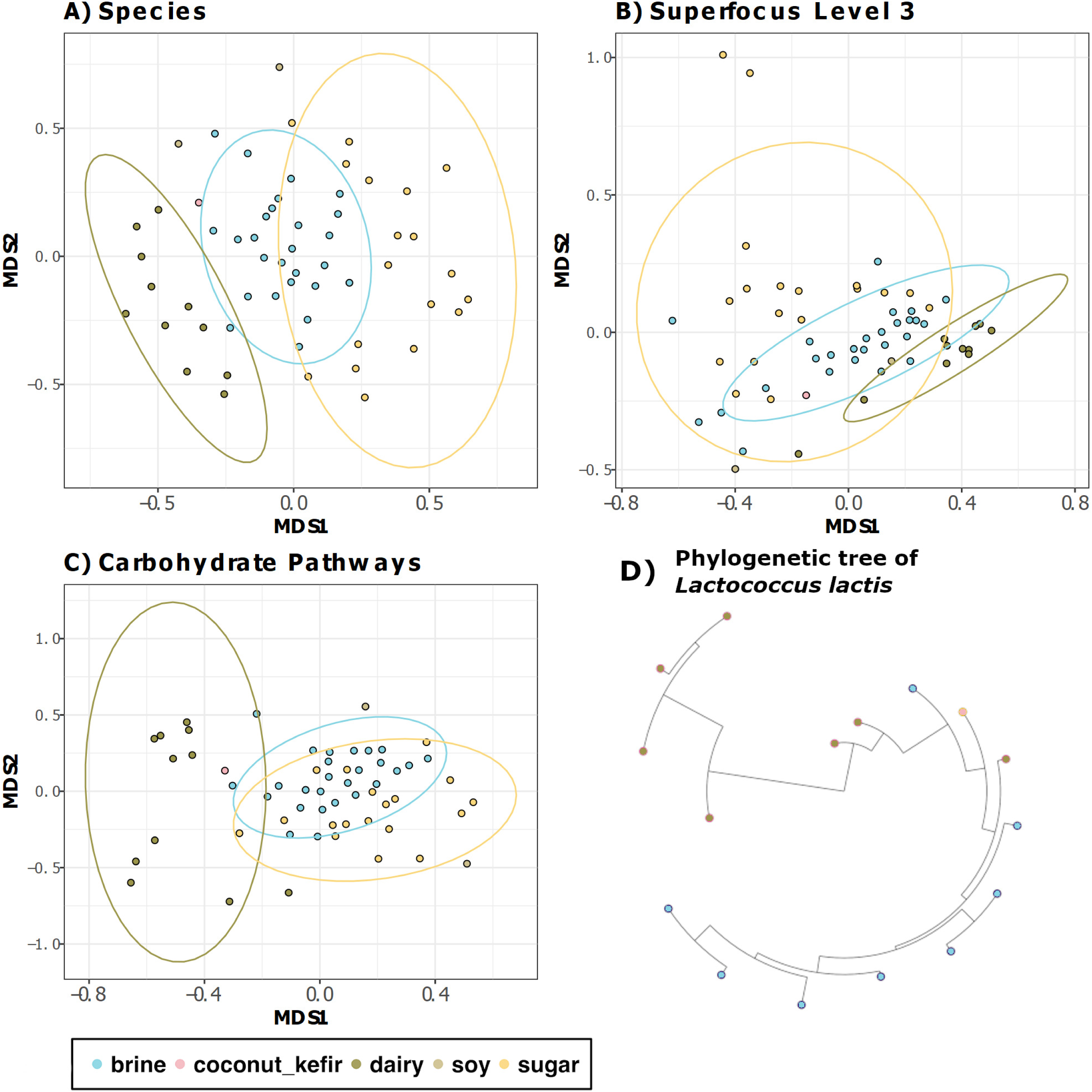
Beta diversity. **A)** Non-metric Multidimensional Scaling (NMDS) of Bray-Curtis distances between the 58 samples, calculated for the species level composition. Samples are coloured by substrate **B)** NMDS of Bray-Curtis distances between the 58 samples, calculated for the Superfocus Level 3 composition. Samples are coloured by substrate **C)** NMDS of Bray-Curtis distances of Carbohydrate pathways assigned with Humann2. Samples are coloured by substrate **D)** Maximum likelihood phylogenetic tree of 16 Lactococcus lactis strains from different food samples. Strains are coloured according to food substrate source. All figures show clear shifts in samples/strains by substrate.

**Table 1:** Table of Fermented foods and metadata. Origin is country of origin, Producer is a numeric code for each producer whom supplied foods, Substrate is the main ingredient fermented, State discriminates between solid and liquid foods and Fermentation refers to whether a starter culture was used or not.

| Sample | ID | Origin | Producer | Substrate | State | Fermentation |
| --- | --- | --- | --- | --- | --- | --- |
| Wagashi Rind | FS00a | Benin | 1 | dairy | solid | Starter |
| Wagashi Core | FS00b | Benin | 1 | dairy | solid | Starter |
| Bread Kvass | FS01 | Russia | 2 | sugar | liquid | Starter |
| Carrot Kimchi | FS02 | UK | 2 | brine | solid | Spontaneous |
| Boza | FS03 | UK | 2 | sugar | liquid | Starter |
| Turnip | FS05 | UK | 2 | brine | solid | Spontaneous |
| Orange | FS06 | UK | 2 | sugar | solid | Spontaneous |
| Krauthehi (sauerkraut) | FS07 | Germany | 2 | brine | solid | Spontaneous |
| Tepache | FS08 | Mexico | 2 | sugar | liquid | Spontaneous |
| Ginger Beer | FS09 | UK | 2 | sugar | liquid | Spontaneous |
| Tempeh | FS10 | UK | 2 | soy | solid | Starter |
| Cucumber | FS11 | UK | 2 | brine | solid | Spontaneous |
| Milk Kefir | FS12 | UK | 2 | dairy | liquid | Starter |
| Water Kefir | FS13 | UK | 2 | sugar | liquid | Starter |
| Tofu Chilli | FS16 | China | 3 | soy | solid | Spontaneous |
| Daikon | FS17 | China | 3 | brine | solid | Spontaneous |
| Pickled vegetables | FS19 | China | 3 | brine | solid | Spontaneous |
| Raw Sauerkraut & Juniper berries | FS22 | Ireland | 4 | brine | solid | Spontaneous |
| Brown rice amazake | FS23 | Japan | 4 | brine | solid | Spontaneous |
| Beetroot Kvass | FS24 | Ireland | 5 | brine | liquid | Starter |
| Kefir and fennel soup | FS25 | Ireland | 5 | dairy | liquid | Starter |
| Mead | FS26 | Ireland | 5 | sugar | liquid | Spontaneous |
| Sauerkraut | FS27 | Ireland | 5 | brine | solid | Spontaneous |
| Dill dearg (sauerkraut) | FS28 | Ireland | 6 | brine | solid | Spontaneous |
| Kimchi | FS29 | Ireland | 6 | brine | solid | Spontaneous |
| Golden child (sauerkraut) | FS30 | Ireland | 6 | brine | solid | Spontaneous |
| Water Kefir Hibiscus | FS31 | Ireland | 6 | sugar | liquid | Starter |
| Water Kefir lemon | FS32 | Ireland | 6 | sugar | liquid | Starter |
| Water Kefir Ginger | FS33 | Ireland | 6 | sugar | liquid | Starter |
| Kombucha Vinegar | FS34 | Ireland | 6 | sugar | liquid | Starter |
| RYAZHENKA | FS35 | Russia | 7 | dairy | liquid | Starter |
| Agousha | FS36 | Russia | 7 | dairy | liquid | Starter |
| ROSTAGROÈKPORT VOROŽNyJ | FS37 | Russia | 7 | dairy | solid | Starter |
| RUŽ'A | FS38 | Russia | 7 | dairy | solid | Starter |
| Sauerkraut | FS39 | Ireland | 8 | brine | solid | Spontaneous |
| Kombucha | FS40 | Ireland | 8 | sugar | liquid | Starter |
| Apple Cider Vinegar | FS41 | Ireland | 8 | sugar | liquid | Starter |
| Raw Milk Kefir | FS42 | Ireland | 9 | dairy | liquid | Starter |
| Pasterised Milk Kefir | FS43 | Ireland | 9 | dairy | liquid | Starter |
| Water Kefir(Pear, Ginger & Honey) | FS44 | Ireland | 9 | sugar | liquid | Starter |
| Water Kefir(Pear, Ginger & Sugar) | FS45 | Ireland | 9 | sugar | liquid | Starter |
| Dilly Carrots | FS46 | Ireland | 10 | brine | solid | Spontaneous |
| Brussel Sprout Kimchi | FS47 | Ireland | 10 | brine | solid | Spontaneous |
| Kimchi | FS48 | Ireland | 10 | brine | solid | Spontaneous |
| Garlic Kraut | FS49 | Ireland | 10 | brine | solid | Spontaneous |
| Dukkah Kraut | FS50 | Ireland | 10 | brine | solid | Spontaneous |
| Ginger Sliced in 2% Brine | FS51 | Ireland | 10 | brine | solid | Spontaneous |
| Daikon Radish in 2% Brine | FS52 | Ireland | 10 | brine | solid | Spontaneous |
| Okra in 2% Brine | FS53 | Ireland | 10 | brine | solid | Spontaneous |
| Tomatoes & mustard seeds in |  |  |  |  |  |  |
| 2% Brine | FS54 | Ireland | 10 | brine | solid | Spontaneous |
| Kombucha | FS55 | Ireland | 10 | sugar | liquid | Starter |
| Cherry Water Kefir | FS56 | Ireland | 10 | sugar | liquid | Starter |
| Beet Kvass | FS57 | Ireland | 10 | brine | liquid | Starter |
| Coconut Kefir | FS58 | Ireland | 5 | coconut_kefir | liquid | Starter |
| Carrot sticks | FS59 | Ireland | 5 | brine | solid | Spontaneous |
| Labne | FS60 | Ireland | 5 | dairy | solid | Starter |
| Lemon and Ginger Fizz | FS61 | Ireland | 5 | sugar | liquid | Starter |
| Scallion Kimchi | FS62 | Ireland | 5 | brine | solid | Spontaneous |

**Table 2:** Anosim results order by descending R statistic. Only results that remained significant (p < 0.05) after Benjamini-Hochberg (BH.padj) corrections are included here (full table **Supplementary** Table 6).

| Level | Variable | P.Value | R.statistic | BH.padj |
| --- | --- | --- | --- | --- |
| Family | Type | 0.001 | 0.651 | 0.008 |
| Genus | Type | 0.001 | 0.551 | 0.013 |
| Carbs | Type | 0.001 | 0.514 | 0.004 |
| Species | Type | 0.001 | 0.436 | 0.050 |
| Superfocus Level 3 | Type | 0.001 | 0.345 | 0.004 |
| Superfocus Level 1 | Type | 0.001 | 0.289 | 0.005 |
| Phylum | Type | 0.001 | 0.280 | 0.006 |
| Carbs | Producer | 0.001 | 0.221 | 0.004 |
| Superfocus Level 2 | Type | 0.001 | 0.210 | 0.005 |
| Family | Fermentation | 0.001 | 0.202 | 0.006 |
| Species | Fermentation | 0.001 | 0.171 | 0.017 |
| Species | State | 0.001 | 0.169 | 0.025 |
| Family | State | 0.001 | 0.167 | 0.007 |
| AMR | Type | 0.004 | 0.163 | 0.010 |
| Species | Producer | 0.003 | 0.160 | 0.008 |
| Carbs | Fermentation | 0.001 | 0.154 | 0.003 |
| Genus | Fermentation | 0.001 | 0.149 | 0.010 |
| Superfocus Level 1 | State | 0.002 | 0.117 | 0.006 |
| Superfocus Level 3 | Fermentation | 0.002 | 0.111 | 0.006 |
| AMR | Fermentation | 0.005 | 0.106 | 0.012 |
| Genus | State | 0.007 | 0.097 | 0.015 |
| Superfocus Level 3 | State | 0.006 | 0.094 | 0.013 |
| Superfocus Level 1 | Fermentation | 0.002 | 0.093 | 0.006 |
| Superfocus Level 2 | Fermentation | 0.006 | 0.080 | 0.014 |
| Superfocus Level 2 | State | 0.012 | 0.076 | 0.024 |
| Carbs | State | 0.019 | 0.073 | 0.035 |
| Bacteriocin | State | 0.018 | 0.070 | 0.035 |

Taxonomy was the most distinguishing feature of the food substrates, as measured by the R statistic, supported by NMDS plots, PLS-DA and *L. lactis* phylogenetic tree (**Figure 1**, **Figure 2**, **Table 2**). Substrate-related differences were greatest at the family-level, but were also significant at the species, genus and phylum level (**Table 2**). Further analysis was implemented at strain level. Examination *of Lactococcus lactis*, the species present across the greatest number of food samples revealed that strains phylogenetically cluster according to food substrate (**Figure 1**). There was no clustering of *L. lactis* strains according to any other factor. Functional analysis revealed that substrate had the most considerable impact on the functional profile of the foods (**Table 2**, **Figure 1**). Carbohydrate pathways most considerably differed across the food groups (**Table 2**). Indeed, of the features examined, the bacteriocin profile was the only characteristic that was not statistically different across the food substrates.

**Figure 2.**
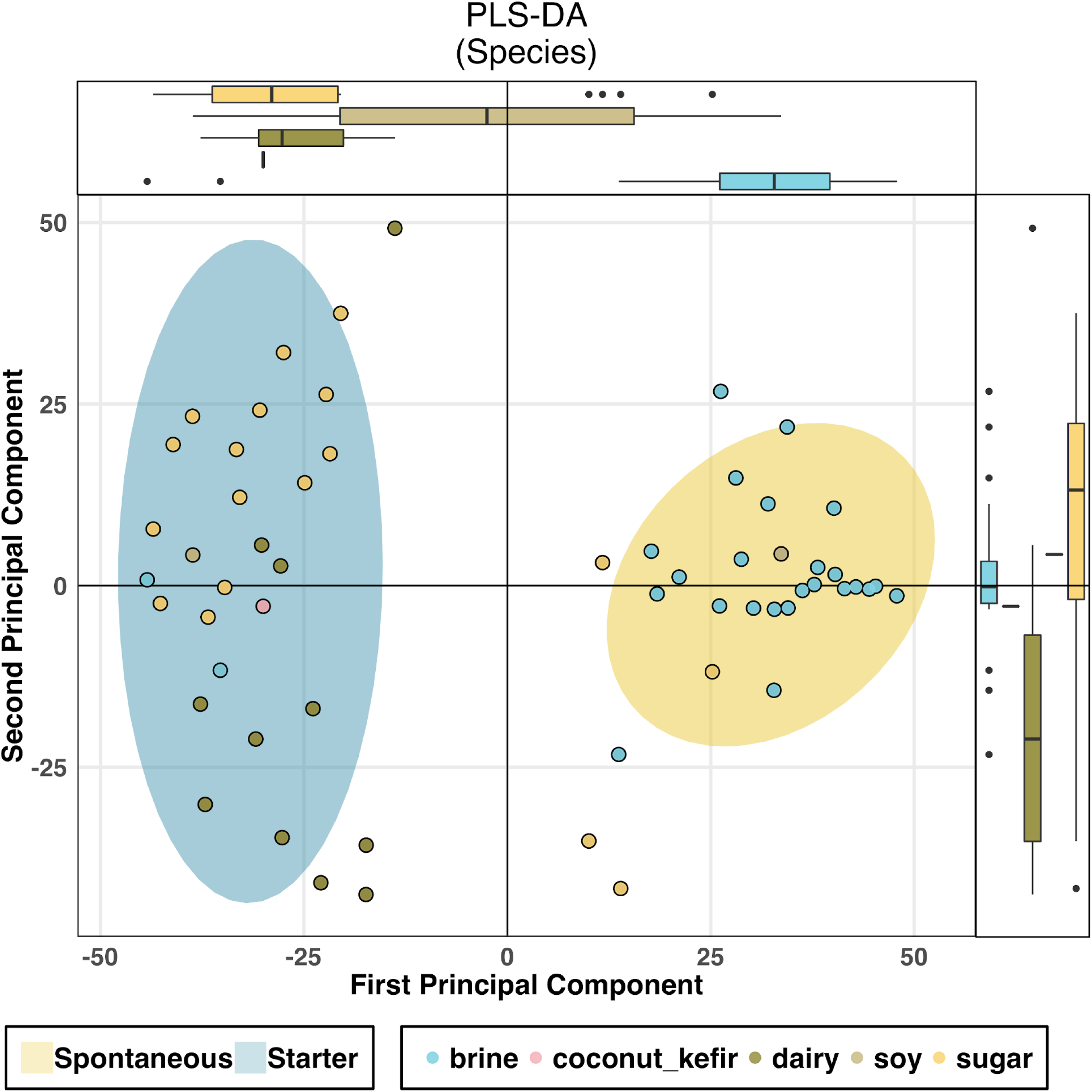
PLS-DA. Variance of sample clustering according to fermentation process and primary substrate. PLS-DA constrained ordination of samples according to fermentation process, illustrates that not all samples exhibit coordination of detected species composition that is dependent on the classification of fermentation process. Samples deviating from the core fermentation-type clusters show unique compositions. PLS-DA, Partial least squares discriminant analysis. Ellipses represent confidence levels of 0.9 of the respective data. Axis plots are boxplots of the plotted data, illustrating distribution of samples according to axis.

Three foods tested did not correspond to the three main food substrates or the corresponding microbiome clusters. Two of these were derived from soy-based fermentations, which are known for their alkaline fermentation environment[50], and the third was a coconut kefir, i.e., a dairy kefir grain based fermentation but of a coconut carbohydrate. Other fermented food types, e.g. fermented meats and fish, were not considered for this study.

### Starter presence/absence, solid/liquid state and producer contribute to differences in microbiota

Although less obvious from a clustering perspective, other factors such as starter presence/absence, solid/liquid state and producer, were also significant drivers of microbiome differences (**Supplementary Figure 1, Table 2**). The presence or absence of a starter culture was associated with differences in family, species, carbohydrate, genus, SF3 and the AMR profile of foods (in order of descending effect size), but to a lesser extent than substrate. Solid/liquid state was significant at three taxonomic levels and all 4 functional profiles (3 SuperFocus levels and Humann2 carbohydrate pathways), but again with a smaller effect size than substrate and starter status (**Table 2**). However, it was the only factor that was associated with significant differences across bacteriocin profiles. The specific producer of the foods was reflected by the carbohydrate related functions and species composition, but country of origin did not influence any of the factors investigated (**Table 2**).

### Microbial diversity differs between dairy foods and other food types

Overall, 476 unique species, present at above 0.1% relative abundance, were assigned to the 58 foods, whereof 301 different species were detected in brine foods, 242 in sugar foods and 70 in dairy. This corresponded to an average of 11.5, 13.5 and 6.4 different species per sample for brine, sugar and dairy foods, respectively. In line with these results, alpha diversity analyses demonstrated that the microbiomes of dairy-based fermented foods had significantly lower alpha diversity than those of either brine or sugar foods (**Figure 3**), which did not significantly differ from one another. It was also evident that, as expected, the alpha diversity of spontaneously fermented foods was significantly higher than those produced using starter cultures (**Figure 4**). Across the specific foods, a spontaneously fermented orange preserve contained the highest number of species (67), while a sample of tepache, a slightly alcoholic spontaneously fermented drink from Mexico, contained the lowest number of observed species (12).

**Figure 3.**
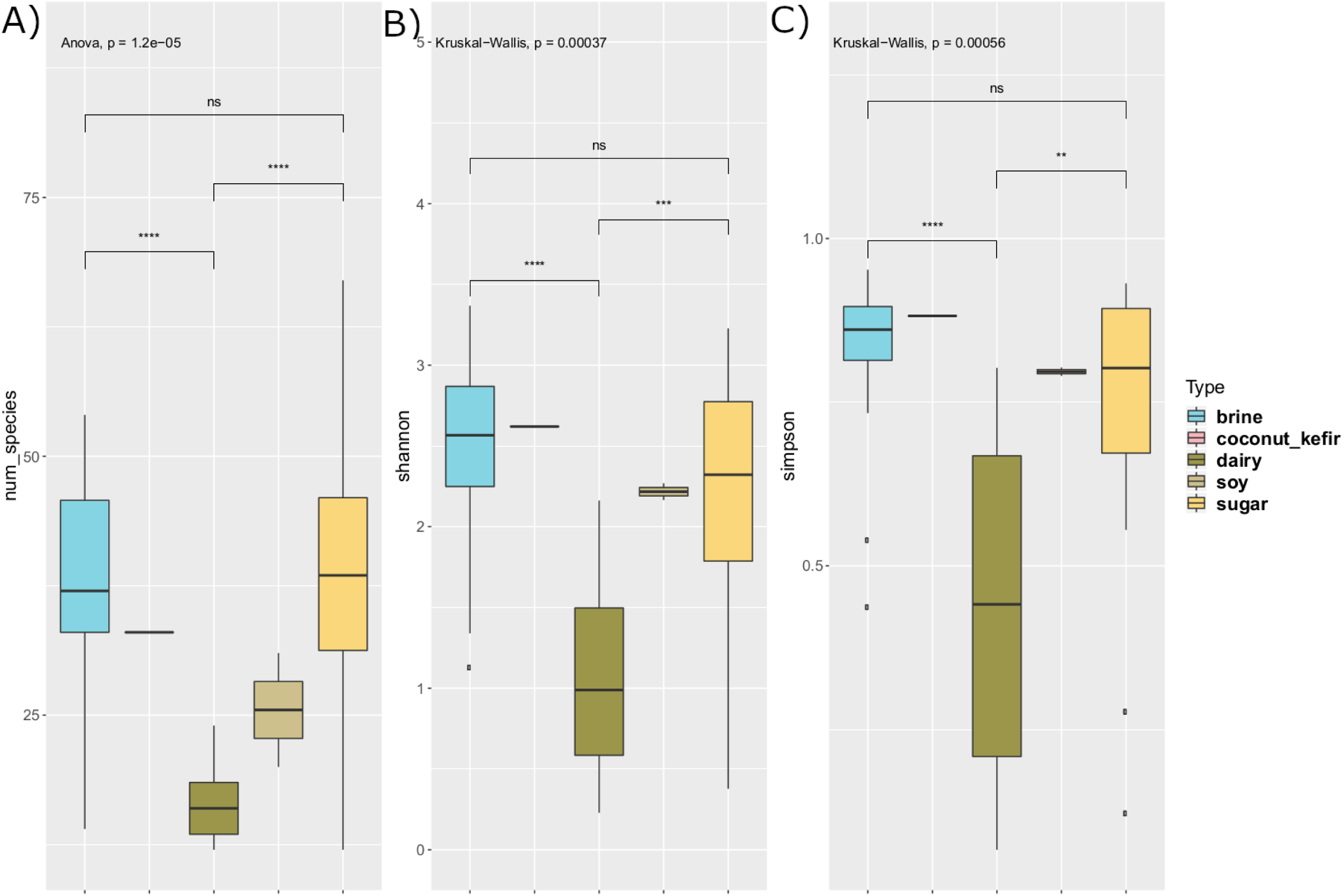
Alpha diversity by substrate. **A)** Number of species (abundance higher than 0.1%) per sample. Anova was used as the data had a normal distribution. **B)** Shannon index of samples. Kruskal-Wallis was used as the data was non-parametric. **C)** Simpsons diversity index of samples. Kruskal-Wallis was used as the data was non-parametric. In all three, the pairwise tests were carried out between Dairy, Brine & Sugar (T-test for parametric and Wilcoxon pairwise test for non-parametric). Coconut kefir and Soy had insufficient sample size for pairwise comparisons

**Figure 4.**
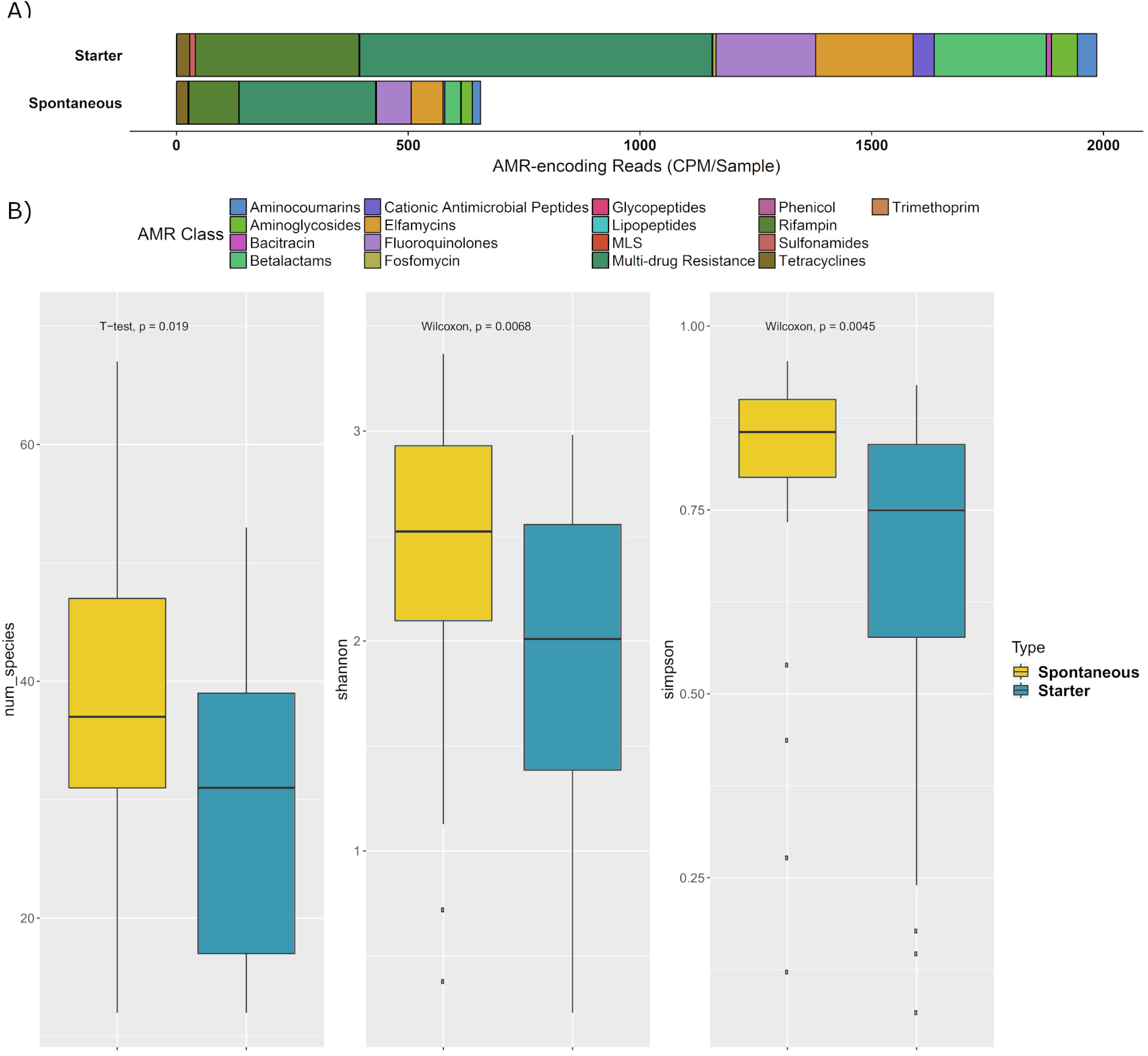
Differences by Fermentation. **A)** AMR profile of Spontaneous fermented foods and Starter culture foods. The AMR classes are normalised by counts per million per sample (CPM). **B)** Alpha diversity boxplots examined across Fermentation type (Spontaneous or Starter). T-test was used for number of species as data was parametric, Wilcoxon test was used for Shannons diversity index and Simpsons index as data was non-parametric.

### Lactic Acid Bacteria dominate Brine Foods

The brine-type foods tested comprised 26 plant substrate-derived foods fermented in a saline solution. Unlike both dairy- and sugar-type fermented foods, the majority of the brine-based foods undergo a spontaneous fermentation and, therefore, rely on fermentation by autochthonous microbes[51]. Among brine-type foods, *Lactobacillus* was the most abundant genus, comprising 46.8% of all reads assigned at the genus level. *Lactobacillus plantarum* was the most abundant species (9.6% relative abundance on average) followed by *L. brevis* (7.9%)*, L. mucosae* (4.7%)*, L. xianfangensis* (4.1%) and *L. sakei* (3%). *Leuconostoc mesenteroides* (4.7%) and *Pediococcus parvulus* (4.3%) were also present in significant quantities. Across the brine-type foods *Bifidobacteriaceae* were detected at a relative abundance of 1.6%. At the species level, 0.8% of species were assigned as *Bifidobacterium longum* and 0.01% *to B. breve*. No other bifidobacteria were assigned at the species level.

Several brine fermented foods were described for the first time, alongside foods that have been described before. A detailed description of these foods can be found in the supplementary material. From a functional potential perspective, 18.4% of Superfocus level 1 (SF1) functions within the brine food microbiome were predicted to relate to carbohydrate metabolism. When functional pathways were investigated at a deeper level, xylose utilisation (0.6%, SF3), fermentation (1.4%, SF2) and response osmotic stress (1%, SF2) were among the most common functionalities (**Supplementary Table 2**). A complete list of the relative abundances of the SuperFocus pathways, for all foods, can be found in **Supplementary Table 2**.

### The microbiota composition of dairy foods is more homogeneous than that of other fermented foods

Eleven dairy-type fermented foods were studied. Information supplied by the producers established that all of these foods were produced through the use of starter cultures to initiate fermentation, thus likely contributing to their reduced diversity relative to other foods [21]. *Lactococcus lactis* dominated, corresponding to, on average, 44.8% of relative abundance and was present at a relative abundance at or above 90% in 3 of the dairy foods, all of which were kefir or kefir-type foods. The next most abundant species was *Streptococcus thermophilus* (16%), followed by *S. infantarius* (5.7%), *Kluyveromyces marxianus* (3.7%), *Escherichia coli* (3.5%), *Lactococcus raffinolactis* (3%) and *L*. *mesenteroides* (2.9%). It was notable that viruses (including (pro)phage) also made up a significant portion of the dairy food microbiota (7.8%). Specific microbiomes of the various dairy fermented foods are described in the supplementary material.

At a functional level, carbohydrate metabolism (16.7%) was the most abundant SF1 pathway in fermented dairy. SF2 results highlighted the presence of genes with homology to those encoding resistance to antibiotics and the production of toxic compounds (2.8% of the reads). Several of the most abundant SF3 pathways in dairy foods had phage related functions, including the most abundant function, i.e., phage head and packaging (3.2%).

### Sugar foods are dominated by *Acetobacteraceae*

Eighteen sugar-type fermented foods were assessed, including fermented fruit, kombucha and water kefir. Some of these foods, such as kombucha, kvass and water kefir, contained large quantities of added table sugar, whereas the substrates used for the production of fermented orange or mead, honey and water, had naturally high levels of sugar. Furthermore, while these foods were all assigned to the ‘sugar foods’ category (**Table 1**), they encompassed a wide variety of raw ingredients and fermentation methods, including examples of both spontaneous and starter type fermentations.

Sugar foods contained many species previously associated with alcohol-generating fermentations, such as *Saccharomyces eubayanus* (2.7%)*, Brettanomyces bruxellensis* (5.2%)*, Hanseniaspora valbyensis* (9.3%) and *Oenococcus oeni* (5%). Many of the other species were well-known kombucha-associated species such as *Gluconobacter oxydans* (5%), *Acetobacter cerevisiae* (2.5%) and *Komagataeibacter rhaeticus* (2%). At the species level, *Hanseniaspora valbyensis* was the most abundant (9.3% average abundance). However, this reflects very high abundance in specific instances, e.g., relative abundance in mead was 93.7%, whereas this species was not detected in 10 of the other 18 sugar-type fermented foods. *Lactobacillus* was the most abundant genus (25.8%) but its abundance was lower than that found for dairy and brine foods. Within this genus, *Lactobacillus mali* (7.6%) and *L. plantarum* (5.3%) were the most common species. *Acetobacter* was the next most abundant genus (10.9%) and its distribution, along with other members of the *Acetobacteraceae*, made it the most abundant family (33.3%). Like brine and dairy fermented foods, the specific microbiomes of sugar foods are described in the supplementary material.

The most abundant SF1 function found in sugar foods was carbohydrate metabolism (14.5%). Resistance to antibiotics and toxic compounds (3.8%) and osmotic stress (1%) were the most common SF2 functions, while analysis at SF3 pathways highlighted the frequency of several pathways involved in the synthesis of amino acids such as methionine (0.79%), as well as purine (0.68%) biosynthesis.

### The fermented food resistome differs according to food and fermentation type

Large variability in both the counts per million of antimicrobial genes (CPM) and of antimicrobial resistance (AMR) class were apparent across the different foods, with AMR profiles significantly differing across substrate and in line with the presence/absence of a starter (**Figure 4**, **Figure 5D**, **Table 2**). Dairy had an average of 3686 CPM per sample, brine had 426 CPM and sugar had 261 CPM. However, the core and the rind of wagashi inflated the dairy results and, if these are excluded, the average CPM for dairy foods dropped considerably to 1947.

**Figure 5.**
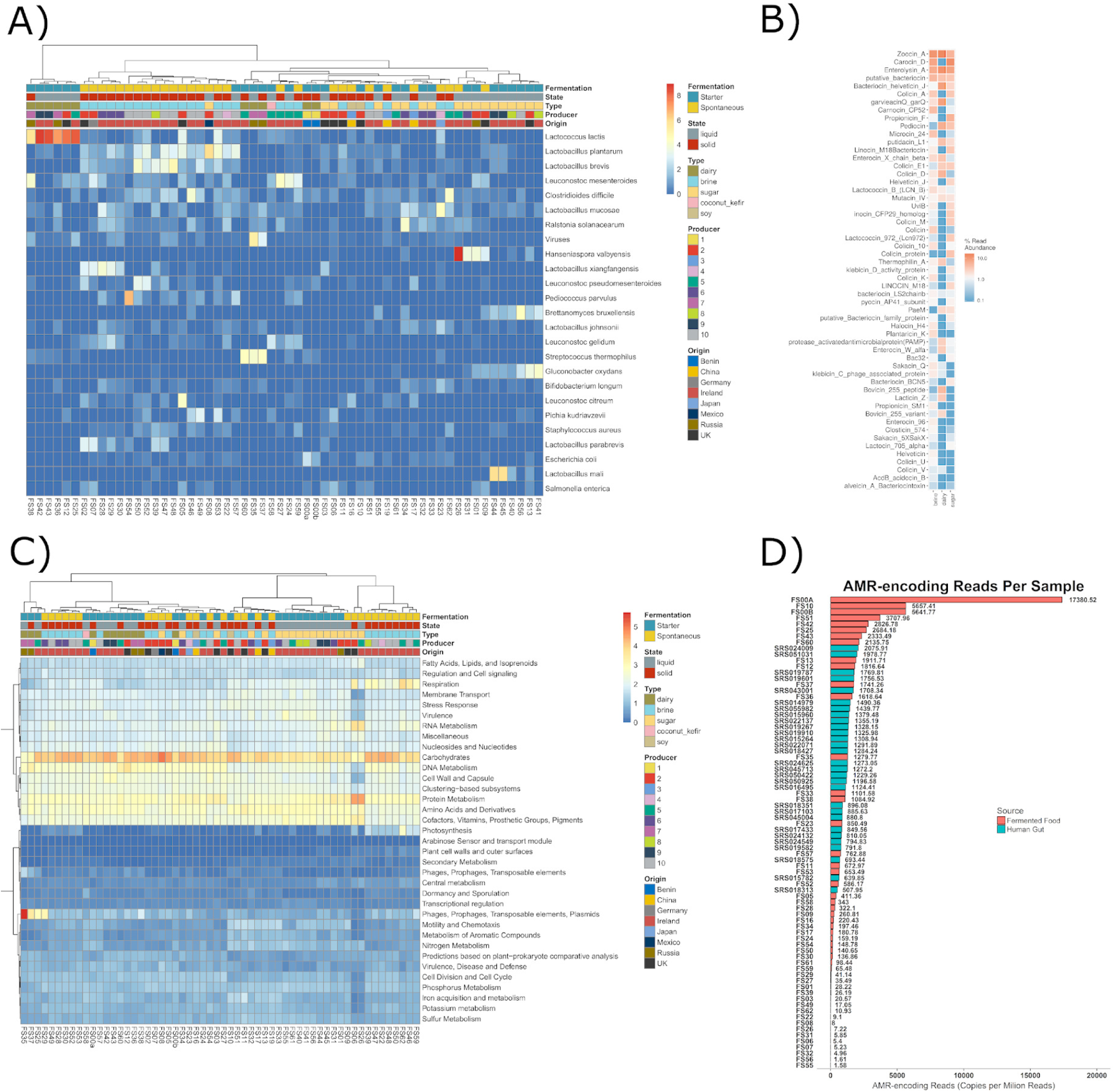
Descriptive plots. **A)** Heatmap showing the square root of the relative abundance of the top 50 Species across all foods. Metadata categories along the top x-axis. Both rows and columns are clustered according to similarity. **B)** Heatmap showing the relative abundance of the bacteriocin profile binned according to food substrate. **C)** Heatmap showing the square root of the relative abundance of the SuperFocus level 1 pathways **D)** Anti-microbial resistance (AMR) genes in counts per million (CPM) per food (pink) and per human sample (blue).

With respect to specific AMR classes, multi-drug resistance was most commonly assigned gene category across all three food substrates, corresponding to 2422, 293 and 133 CPMs per sample on average for dairy, brine and sugar-type foods, respectively. Betalactam resistance genes were the next most common class in dairy (718 CPM) and sugar (101 CPM) foods, while tetracycline resistance genes were the second most numerous category of AMR genes in brine (45 CPM). It was also noted that a five-fold higher abundance of AMR genes occurred in starter culture fermentations relative to spontaneous fermentations. Multi-drug resistance genes again dominated, corresponding to 1326 CPM for starter cultures and 236 CPM for spontaneous fermentations. Betalactam resistance genes were the next highest in foods containing starter cultures (428 CPM), whereas tetracycline resistance genes were next highest in spontaneously fermented foods (48 CPM). The high CPMs for both dairy and starter containing foods is consistent with the fact that dairy foods were those for which starters were most extensively used. When gene distribution was investigated from the perspective of specific food substrates, the wagashi cheese rind was found to have the highest CPM, i.e., 17381, with tempeh being next highest with 5657 CPM. AMR genes counts in kombucha and water kefirs were generally low, and no known AMR genes were identified 9 of the 58 foods, i.e., 1 kombucha, 2 water kefirs, 3 kimchi, 1 pickled carrot, 1 pickled vegetable and 1 apple cider vinegar. Of the 9 fermented foods for which no AMR genes were assigned, 4 were sugar-type (including 2 water kefirs) and 5 were brine-type (including 3 kimchis). It was notable that very few AMR genes were assigned in the 2 other kimchis studied (<42 CPM) while across the 5 other water kefir samples, 3 contained very few AMR genes (<6 CPM) but 2 had relatively high counts (>1000 CPM). Across the two samples of Kombucha, 1 did not contain assigned AMR genes while the other contained 1.6 CPM.

To provide context, the frequency with which AMR genes are detected in fermented foods was compared with that across human stool samples for comparative purposes (**Figure 5D**). Human gut samples (29 random stool samples from the Human Gut Microbiome Project[52]) had significantly more AMR CPMs than fermented foods (**p > 0.01**) with the exception of 8 fermented foods. These 8 foods were the 2 wagashi cheese samples, tempeh, fermented ginger, 3 milk kefirs and labne. Of these 8 foods, 6 were dairy, and 7 were starter-generated foods. A further 12 foods had similar CPM of AMR genes, while 38 foods had lower AMR CPMs, when compared with the human samples.

### The presence of putative health promoting genes differs markedly across fermented foods but exceeds that of non-fermented foods

Bacteriocins are ribosomally synthesised antimicrobial peptides, many producers of which have been sourced from fermented foods. The bacteriocin-producing potential across the 58 fermented food samples was investigated, with 55 putative bacteriocin-encoding gene clusters being assigned across 54 of the foods (no gene clusters identified in 4 samples (**Supplementary Table 3**). Zoocin A- and enterolysin A-like gene clusters were highly abundant across all 3 fermented food substrates.

Clusters corresponding to another bacteriolysin subclass, the helveticin J-like proteins were more frequently detected in dairy and sugar-type foods than in brine-type foods (**Fig 3B**). Carocin A- and colicin A-like clusters had a high abundance in brine and sugar, but not dairy, foods. As noted above, there was a significant difference in the distribution of bacteriocins between solid and liquid food types (**Table 2**), with liquid foods having a higher relative abundance of helveticin J Propioncin F-like and pediocin clusters and solid foods having more carnocin CP52-like and microsin 24-like clusters. Examining the pediocin sequences in more detail, homology with *pedA* and *pedB* was discovered.

Given that bacteriocin production is regarded as a probiotic trait, these findings prompted an investigation of other potentially health-associated gene clusters (PHAGCs) within these fermented food microbiomes. PHAGCs were divided into 3 broad categories. Gene clusters binned as “survival” are genes that were shown to be important for surviving the low pH of the stomach or the bile salts of the small intestine [53]. Gene clusters binned as “colonisation” are genes which were shown to be vital for colonising the gut microbiome. These included genes responsible for surface proteins and exopolysaccharide production. “Modulation” gene clusters were all of the other potentially health promoting gene clusters that did not fit the previous two bins. These genes were shown to affect the host phenotype in other ways, such as stimulating the host immune system in the case of D phenyllactic acid [13] or the production of γ aminobutyric acid (GABA) [54, 55]. The majority of these PHAGCs genes are based on studies reviewed in [53]. Shotgun metagenomic data from non-fermented foods, i.e., unpasteurized whole milk, pasteurized skimmed milk and milk powder, was used for comparative purposes. In general, the fermented foods contained considerably more PHAGCs than the non-fermented substrates. Among the fermented foods, a larger number of PHAGCs were found in brine- and sugar-foods than in dairy foods, with several water kefirs, sauerkrauts, beet kvasses and one kombucha being the foods with highest levels of PHAGCs (**Figure 6**). With respect to the individual PHAGC sub-categories, all fermented foods contained more colonisation-type PHAGCs than the non-fermented controls. In the case of the modulation and survival clusters, the number of PHAGCs in some fermented foods, such as scallion kimchi, labne, agousha and mead, were no greater than those in the non-fermented foods.

**Figure 6.**
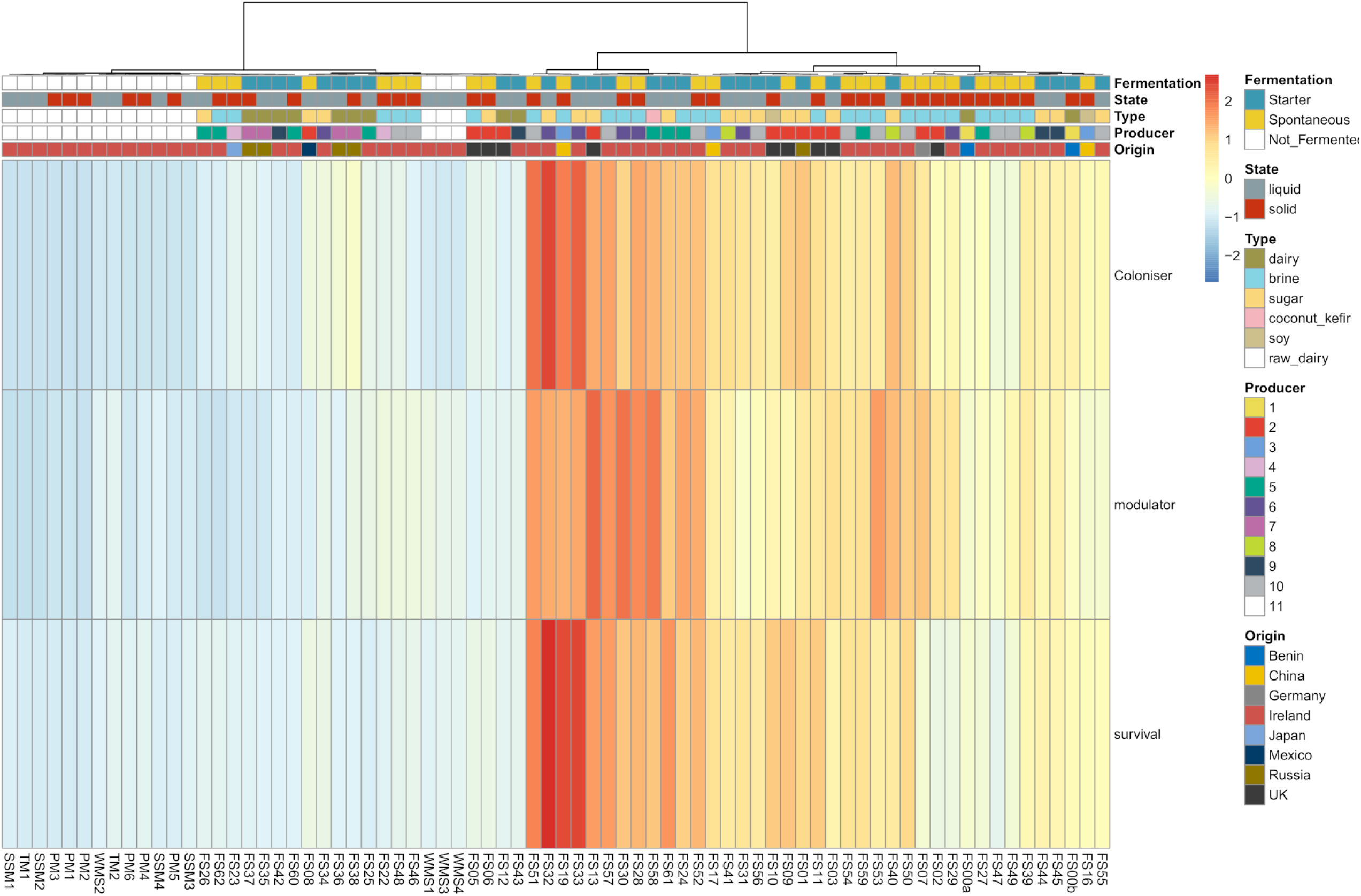
PHAGC heatmap. Heatmap showing the presence ofPotentially Health Associated Gene Clusters (PHAGC) across all 58 foods and 16 unfermented milk samples. Gene clusters are binned as potentially inferring an ability of the metagenome to colonise the gastro-intestinal tract, survive transit to the gut and modulate the host phenotype. Each row is normalised across all samples, therefore only comparing foods to one another.

### Metagenomic assembly reveals 10 putative new species

Metagenome assembled genomes (MAGs) were assembled from the reads and quality checked. 443 MAGs were assembled in total, with 127 genomes above 80% completeness and having less than 10% contamination (**Figure 7**). Traitar[56] was used to predict the growth phenotypes of the 127 MAGs. The outputs were concatenated into a single output for each food substrate (**Figure 7**) and provided intuitive results, such as a high correlation between lactose utilisation and dairy foods and high glucose oxidation potential in sugar food microbiomes. Consilience between the Traitar and taxonomic output is supported by the abundance of *Lactococcus lactis* in dairy and brine samples. FastANI[57] was used to assign taxonomy and to assess novelty and established that 10 of these MAGs had <95% identity to known NCBI prokaryote genomes. 7 of these novel MAGs are acetic acid bacteria, 2 are lactic acid bacteria and 1 belongs to the family *Enterobacteriales* (**Table 3**). The highest identity match for 3 of the novel MAGs was *Acidisphaera rubrifaciens.* All 3 of these MAGs came from water kefir. The 4 remaining acetic acid bacteria were best matched with *Acetobacter aceti* (MAG from water kefir), *Gluconobacter cerinus* (MAG from bread kvass) and *Acetobacter malorum* (MAGs from rostagroèkport vorožnyj and apple cider vinegar). The two novel LABs were best matched with *Leuconostoc gelidium* (sauerkraut MAG) and *Lactobacillus kimchiensis* (boza MAG). The final novel MAG, from the water kefir microbiome, most closely resembled *Rouxiella chamberiensis*.

**Figure 7.**
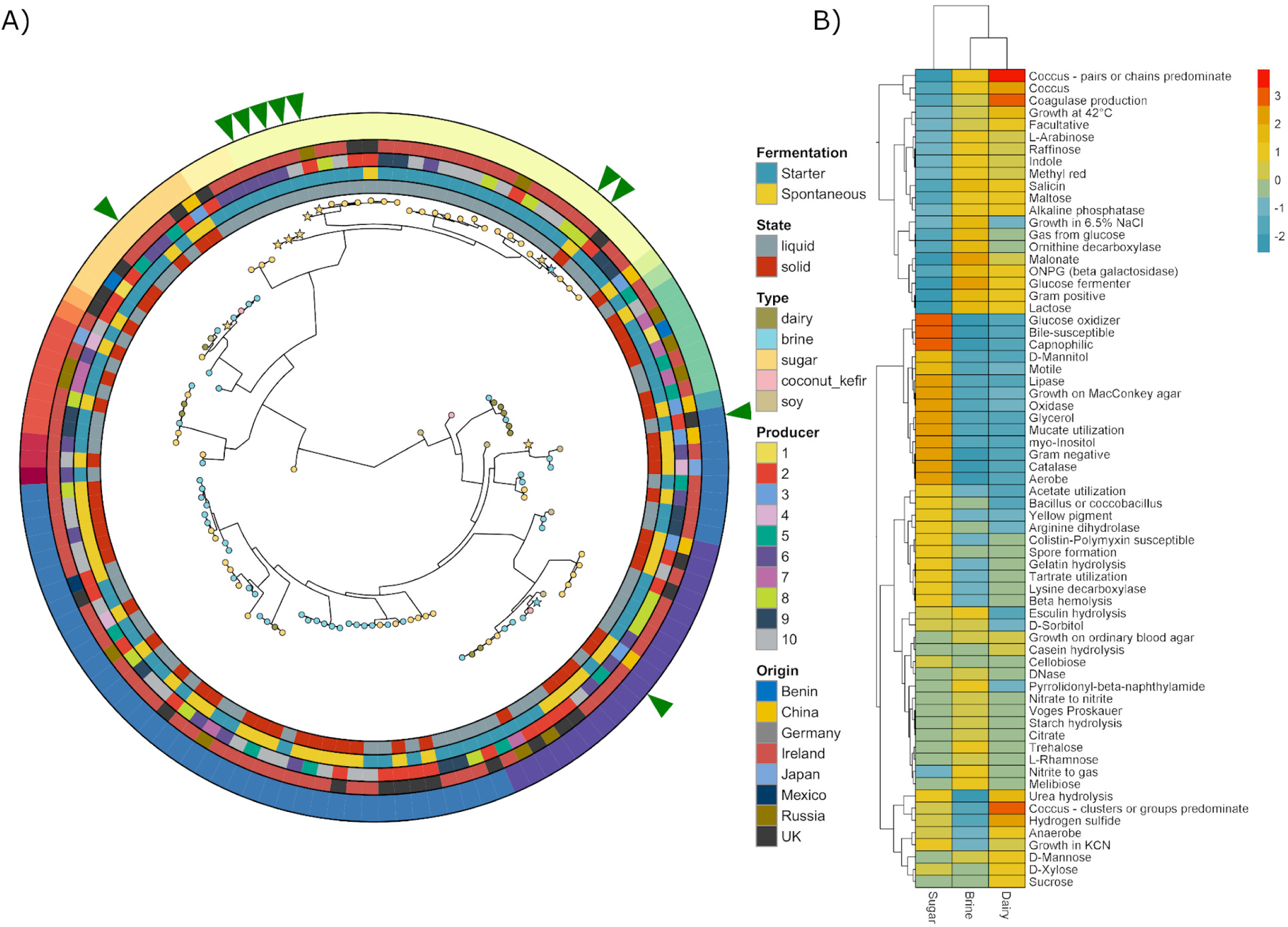
Metagenome Assembled Genomes. **A)** Phylogenetic tree of the 127 high quality MAGs with outer rings showing the metadata of the food. The green arrows indicate which MAGs are potentially novel species. **B)** Predicted phenotypes of the 127 MAGs concatenated into their respective substrate. Both rows and columns are clustered according to similarity.

**Table 3:** Putatively novel MAGs with FastANI identity scores to the closest genome in the NCBI database.

| Food | Sample | Closest NCBI match | Identity |
| --- | --- | --- | --- |
| Bread Kvass | FS01 | Gluconobacter cerinus | 93.4228 |
| Raw Milk Kefir | FS41 | Acetobacter malorum | 86.3852 |
| Sauerkraut | FS39 | Acetobacter malorum | 85.9458 |
| Boza | FS03 | Lactobacillus kimchiensis | 82.2453 |
| Water Kefir lemon | FS32 | Rouxiella chamberiensis | 81.3335 |
| Golden child (Sauerkraut) | FS30 | Leuconostoc gelidum subsp. gasicomitatum | 81.0244 |
| Cherry Water Kefir | FS56 | Acetobacter aceti ATCC 23746 | 78.5186 |
| Water Kefir Hibiscus | FS31 | Acidisphaera rubrifaciens HS-AP3 | 78.4976 |
| Water Kefir Ginger | FS33 | Acidisphaera rubrifaciens HS-AP3 | 78.475 |
| Water Kefir lemon | FS32 | Acidisphaera rubrifaciens HS-AP3 | 78.0727 |

## Discussion

The practice of fermenting foods can be traced back over many millennia[58]. Recently, shifts in consumer preference have resulted in a renewed interest in fermented foods, with the associated global market estimated to reach $40 billion USD by 2024[59]. The development of a better understanding of the microbial composition and functional potential of these foods provides an insight into features that are common among, and different between, fermented foods and ascertain potential roles of individual species, including novel species and strains. Importantly, the taxonomic resolution of shotgun metagenomics allows strain level identification of the microbiome but also facilitates an assessment of functional profile, bacteriocin and AMR gene distribution, determination of PHAGCs, the assembly MAGs and the determination of predicted phenotypes.

Fermentation substrate is the strongest driver of the composition and functional potential of the microbiomes of fermented foods. The type of nutrients available to the microbes determined the diversity within each food to the greatest extent. The biggest effect of substrate was found between the families present in each food substrate, with *Lactobacillaceae* (LDA = 5.68) most persistent in brine foods, *Streptococcaceae* (LDA = 5.92) in dairy foods and *Acetobacteraceae* (LDA = 5.5) in sugar based foods (**Supplementary Figure 2**). The different substrates impose functional requirements on the microbes, such as a necessity for osmotic stress tolerance in both brine and sugar-type foods.

Other factors, such as the presence or absence of a starter culture, also contributed to differences. Starter culture foods had the lowest alpha diversity, likely a result of adding a community of specialist microbes to the food, which would outcompete any autochthonous microbes less adapted to such an environment. Two kefir samples made from the same starter, but using raw or pasteurized milk, respectively, highlight this point. Although we do not have data on the pre-fermented milk, the raw milk likely contained its own unique consortium compared to the relatively low bacterial load of the pasteurised sample. After 48 hours of fermentation, both samples had almost identical microbial composition. The small differences may be due to carry over differences in the microbiota of the substrates, the stochastic differences between any two fermented samples and species falling below the 0.1% abundance threshold for inclusion (hence the appearance of 5 unique species between the 2 samples). Interestingly, *P. helleri* was found at 3% in the pasteurised sample (not at all in the unpasteurised), having been isolated from raw milk in previous studies[60]. The differences in diversity between solid and liquid foods is possibly due to the selective pressures of mobility, nutrient availability (in a homogenous liquid compared to a less homogenous solid food) and moisture content in solid foods compared to liquid foods. The observed differences in diversity due to producer are more difficult to explain, but unrecorded factors such as individual fermentation practises or cross contamination of foods or from the processing environment may be the cause of these differences. Country of origin was not significant for any characteristic examined, possibly due to the cosmopolitan nature of all of the fermenting microbes. Outside of composition and top-level functionalities, other traits did vary in line with other categories, in that bacteriocin gene cluster profile differed significantly across solid and liquid foods, and AMR-encoding genes differed across food substrate and between spontaneous and starter-type fermentations. It is unclear as to why bacteriocin gene clusters differed across solid and liquid foods, but perhaps the matrices of solid foods require different ecological tools for competitive advantage than liquid substrates.

Analysis revealed that the microbiomes of starter culture-type fermentations contain more assigned AMR-associated genes. However, this difference could represent the more extensive characterisation of starter culture microbes, and their associated genomes and AMR profiles, leading to better assignment of AMR genes from starter cultures strains than those involved in spontaneous fermentations. Comparing with human gut metagenomes, the majority of the fermented foods had a lower AMR CPM. Of the 8 foods with higher AMR CPM, only 3 foods stood out as having considerably higher CPMs, 2 were subsamples of the same food, i.e. wagashi cheese. In contrast, kimchi and kombucha samples were notable by virtue of either lacking detectable AMR genes or having very low CPMs. Kimchi shared many taxa with other brine-type foods so the differences observed may reflect strain level differences. Metagenomic sequencing of a larger collection of these fermented foods, coupled with antibiotic resistance assessments of isolated strains, will be necessary to determine how representative these results are.

Bacteriocin production is regarded as a probiotic trait. These peptides and, in the case of bacteriolysins, proteins, are thought to be produced by bacteria to gain a competitive advantage over other taxa, typically those occupying the same environmental niche. Bacteriocin production can contribute to the quality and safety of foods through the removal of spoilage and pathogenic bacteria, but bacteriocin production *in situ* in the gut can also enable the producing bacteria to become established, compete against undesirable taxa and contribute to host-microbe dialogue[61, 62]. The bacteriocin profile did not differ according to food substrate, with zoocin A- and enterolysin A-like genes being most abundant across all food substrates. However, the bacteriocin-associated genes present in solid and liquid foods differed significantly from one another in that liquid foods were enriched with pediocin-like genes. After a further analysis of the pediocin sequences, homology with *pedA* and *pedB*, required for production of to pediocin AcH/PA-1, was apparent. xThese bacteriocins are best known for their strong antilisterial effects[63]. Pediocin AcH/PA-1 has also been shown to be active against enterococci and staphylococci[64], and the presence of these genes potentially adds to the safety of these foods, and their potential to be health promoting. Solid foods had a higher abundance of carnocin CP52-like bacteriocins, which are known for activity against *Listeria* and *Enterococcus*, again potentially adding to the safety of these foods[65].

Across a broader range of PHAGCs, it was apparent that these gene clusters were more common in fermented, than non-fermented, foods. Sugar and brine foods were found to contain the highest levels of PHAGCs. Microbes in sugar-type food microbes generally must persist in low pH environments, with some kombucha fermentations dropping to as low as pH 3[66]. In contrast, although also somewhat acidic, a milk kefir fermentation is regarded as complete when the pH reaches 4.5[67], while the pH of most cheeses is between pH 5.1 and 5.9. Many of the sugar foods also contained colonisation-associated PHAGCs. It was also noted that brine-type foods had the highest abundance of *Lactobacillaceae*, specific representatives of which have been exploited for their probiotic activity. A combination of these various factors likely contributes to the higher abundance of PHAGCs in both of these foods relative to dairy foods. However, even within the respective food substrate groups, the PHAGCs present varied considerably, with foods such as water kefirs, sauerkrauts, pickled veg, ginger, kvass and kombucha being enriched in PHAGCs. These foods all contained colonisation and survival PHAGCs at a higher frequency, e.g., glycotransferases for colonisation in kombucha and pickled veg, and bile salt metabolism genes in water kefir and fermented sliced ginger. D-lactate dehydrogenase pathways were consistently identified in these foods but were absent from other such as scallion kimchi, carrot sticks and agousha. This observation is notable as D-lactate dehydrogenase is the enzyme responsible for producing D-phenyllactic acid (D-PLA), a metabolite known to modulate the host immune system[13]. Glutamate decarboxylase, which converts glutamate into gamma-aminobutyric acid (GABA), was present in some (kombucha, kvass, coconut kefir and some water kefir samples), but not all, PHAGC-enriched foods. GABA is a well-known modulator of mood[68], while this enzymatic reaction also consumes protons and thus contributes to acid resistance[69]. Although *in vivo* studies are required to directly examine the health benefits of specific fermented foods, these insights can undoubtedly help to identify foods, and strains, that are more likely to be health promoting, facilitate the production of fermented foods optimised for health promotion and direct the experimental design of human intervention studies.

Finally, this study discovered 127 high quality MAGs, of which 10 are putative novel species. 3 putative new *Acetobacter* species from water kefir, milk kefir and sauerkraut, a *Gluconobacter* from bread kvass, a *Leuconostoc* from sauerkraut and a *Lactobacillus* from boza were assembled from the shotgun data. While these species are apparently novel, the corresponding genera are found in fermented foods at a high frequency. However, 2 MAGs representing genera that have not been found in fermented foods before were assembled, i.e., a *Rouxiella* species and 3 *Acidisphaera* species, all from water kefir samples. *Rouxiella chamberiensis* and *Acidisphaera rubrifaciens* are the only previously known members of their respective genera. *Rouxiella chamberiensis* was isolated from parenteral nutrition bags and has been shown to ferment D-glucose but not sucrose[70] and *Acidisphaera rubrifaciens* has been found in acidic hot springs and mine drainage systems and, like many of the other sugar taxa, is acidophilic[71]. The assembly of these and other MAGs in the future will contribute towards the building of fermented food, and other food, microbe databases, equivalent to those available for the more complex human gut microbiome[72], to enable the more accurate and rapid identification of food microbes. Such databases will be key in the application of metagenomics-based approaches on a widespread basis by the food industry.

Overall, this study combines many novel insights into fermented food microbiomes. Firstly, the taxonomic composition of the 58 foods has been described, including many foods that have not been described using NGS previously. Secondly, the functional profile of these foods has been characterised, and like the taxonomic profile, highlights the differences between starting material and microbial composition. Importantly, given the current interest in fermented foods as a healthy food choice and the role diet plays in modulating the gutmicrobiome, the health promoting potential of the microbes in these various foods has been explored. Finally, genomes, including potentially novel taxa, were assembled from these foods, and will contribute to the better assignment of reads from fermented food, and indeed broader food chain microbiome studies, in the future.

## Methods

58 samples of fermented foods were collected from various artisanal producers (see Table 1). 5g of solid foods were placed in a stomacher bag. 50ml of sterile MRD was added to the bag. The contents were homogenised in a stomacher (BagMixer 400 from Interscience) for 20 minutes. After this step, both solid and liquid foods were extracted using the same method. 50ml of the homogenised solution was centrifuged at 10,000 rpm, at room temperature, for 10 minutes. The supernatant was discarded. The pellet was resuspended in 550µl of SL buffer in a 2ml tube (SL buffer from GeneAll kit below). 33µl of Proteinase K was added to the tube and incubated at 55°C for 30 minutes. The solution was then transferred to a bead beating tube and placed in a Qiagen Tissue lyser 2 for 10 minutes at 20/s. The GeneAll Exgene extraction protocol from step 4 was then followed until the final elution step, where 30µl of elution buffer (EB) was used here instead of the 50µl suggested in the protocol.

### Sequencing

Library prep was carried out as per Illumina Nextera XT protocol (Illumina) [73]. DNA was quantified using a Qubit High Sensitivity dsDNA assay. Final library quality was assessed by running on an Agilent High Sensitivity DNA chip, and quantification by qPCR using the KAPA Library Quantification Kit for Illumina (Roche). Sequencing was carried out on the NextSeq500 using a 300 cycle High Output v2 kit.

### Bioinformatics

All raw reads can be accessed from the ENA under the project accession number PRJEB35321. 347,841,507 reads were obtained from the Nextseq sequencing run in the form of Bcl files, which were converted to fastq format using bcl2fastq software. Quality trimming was performed using the trimBWAstyle.usingBAM.pl script. Using Picard (https://github.com/broadinstitute/picard), fastq was converted to Sam format. Picard was also used to remove duplicates. The sequences were then quality checked and trimmed using the trimBWAstyle.usingBam.pl script from the Bioinformatics Core at UC Davis Genome Center (https://github.com/genome/genome/blob/master/lib/perl/Genome/Site/TGI/Hmp/HmpSraProcess/trimBWAstyle.usingBam.pl). Forward and reverse reads were then combined into a single fasta file for each sample using the fq2fa command from IDBA-UD [74].

Kaiju [75] was used to assign taxonomy to the reads, discarding taxa with relative abundance of less than 0.1%.This setting was chosen as other studies have shown a high false positive discovery rate below this threshold [76]. Superfocus [77] was used to assign functionality to the reads. All percentages reported at all taxonomic levels are percentages of the assigned reads only. **Supplementary Table 1** shows the complete list of microbes and their relative abundance for each food. The phylogenetic tree of *L. lactis* was created in GraPhlAn [78], using the StrainPhlAn [79] output, which used Metaphlan2 [80] taxonomic assignment.

Statistical analyses was carried out in R-3.2.2[81] using vegan [82]. Anosim analysis was carried out between each metadata category containing 6 or more samples (**Supplementary table 4**). Benjamini-Hochberg false discovery rate was applied to the anosim results. The linear discriminant analysis (LDA) effect size (LEfSe) [83]method was used to determine if any taxa or pathways were differentially abundant between groups.

### Antimicrobial Resistance

Antimicrobial resistome analysis was performed by aligning paired-end metagenomes reads against the MEGAres database (v. 1.0.1) [84]. To reduce Type I errors, this database was first manually curated to remove any genes corresponding to antimicrobial resistance arising from point mutations. The alignment was performed using the --very-sensitive-local preset of Bowtie2 (v. 2.3.4). The Resistome Analyser tool (https://github.com/cdeanj/resistomeanalyzer) was used to format the output and the results were normalised for sequencing depth across samples as copies per million reads (CPM).

### Bacteriocin Assignment

Bacteriocin assignment was performed with the BLAST analysis of the bacteriocin genome mining tool (BAGEL) of the predicted genes with the Prodigal tool against the BAGEL4 bacteriocin databases [85].

### Carbohydrate pathways

The carbohydrate function was assigned to reads with the HUMAnN2 pipeline [86], which assigned the function based on the ChocoPhlan databases and genes based on UniRef [87]. To further simplify the exploration of the abundance data of the gene family were grouped into the functional category Gene Ontology (GO), specifically carbohydrate-related functions, performing a more in-depth analysis.

### Metagenomic Assembled Genomes

Metagenome assembly was carried out using IDBA-UD. MetaBAT 2 [88] was used for genome binning, with default settings. CheckM [89] was implemented to check the quality of metagenome assembled genomes (MAGs). Low quality MAGs, i.e. <80% completeness and/or >10% contamination, were removed from downstream analysis. Kaiju [90]and PhyloPhlAn [91] were used to assign taxonomy to the MAGs. The average nucleotide identity (ANI) of MAGs to reference genomes, which were downloaded from RefSeq [92], was calculated using FastANI [57]. Putatively novel MAGs were assigned as potentially new species using the same ANI threshold as [72]. The phenotypes of MAGs were predicted using Traitar [56]. MAGs were annotated using Prokka [93].

### PLS-DA analyses

Partial least squares discriminant analysis (PLS-DA) plots were generated using the KODAMA R package (version 1.5) [94]. Default parameters of the KODAMA software were used on species from the taxonomic profile with the semi-supervised constraining of data ordination according to the fermentation process of samples. The final visualisation of data was performed in R (version 3.5.1) using ggplot2 (version 3.1.1) [95].

### PHAGC screening

Shotgun sequences for 16 non-fermented dairy samples were downloaded from ENA (study accession number PRJEB31110) with a median of 18041 reads per sample, after removing *Bos taurus* reads. The 16 dairy samples were; raw tanker milk X 2, skimmed milk powder x 6, pasteurized skimmed milk x 4 and raw silo whole milk x 4. The fermented and non-fermented food sequences were then assigned Uniref90 clusters using the Humann2 software[86]. Using the Uniref90 clusters obtained from Humann2 output, the presence or absence of clusters that have been shown to influence potential health promoting properties of bacteria was determined[13, 53, 96]. The list of search terms can be found in **Supplementary table 5**. The total number of PHAGCs present in each food were binned into one of the following 3 categories; survival, modulation and colonisation. The heatmap was created using Pheatmap[97]. The rows of the heatmap were scaled, so that the values are comparative between the foods, and not an absolute count of the number of gene clusters found in each food.

## Acknowledgements

The authors thank the fermented food producers for kindly supplying the samples. This research was conducted with the financial support of Science Foundation Ireland (SFI) under Grant Number SFI/12/RC/2273.

## Author contributions

PDC conceived the study idea and design. JL and GM collected samples and extracted DNA. FC & LF conducted sequencing. JL, RCR, AMW, JCW & WB conducted bioinformatics analysis. PDC and JL wrote the manuscript with contributions from everybody else. OOS, MJC & PDC supervised the project.

---

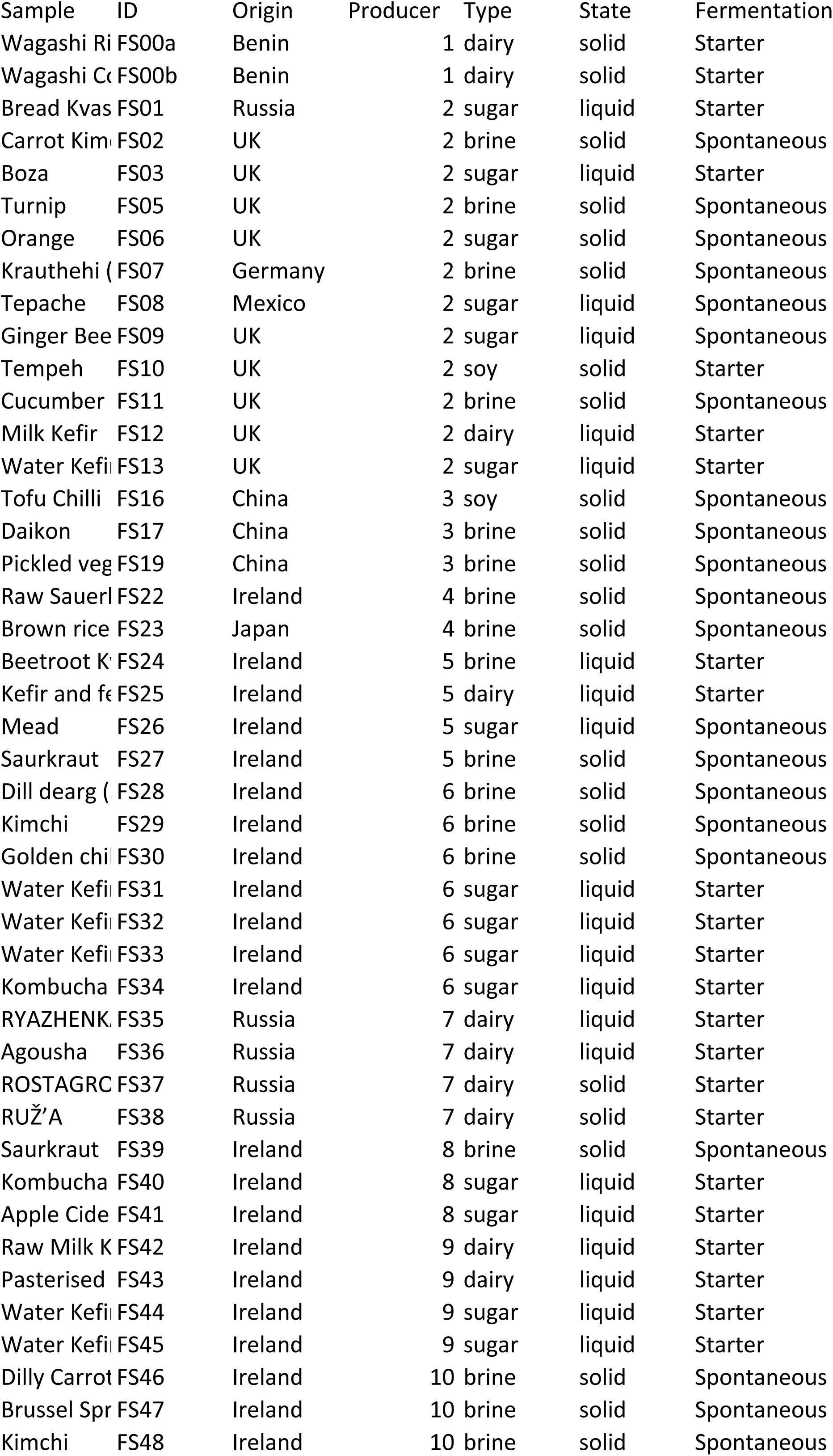
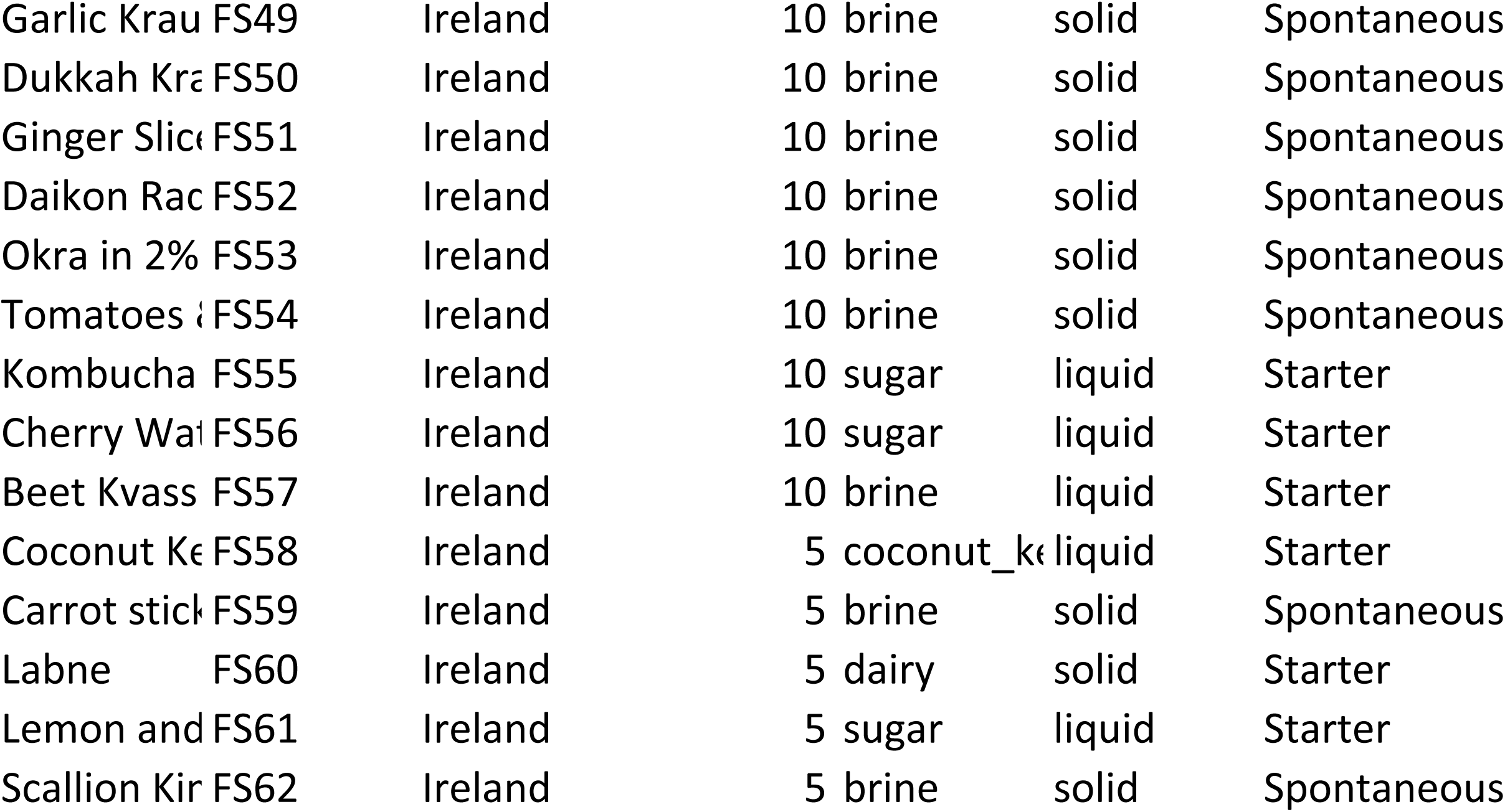
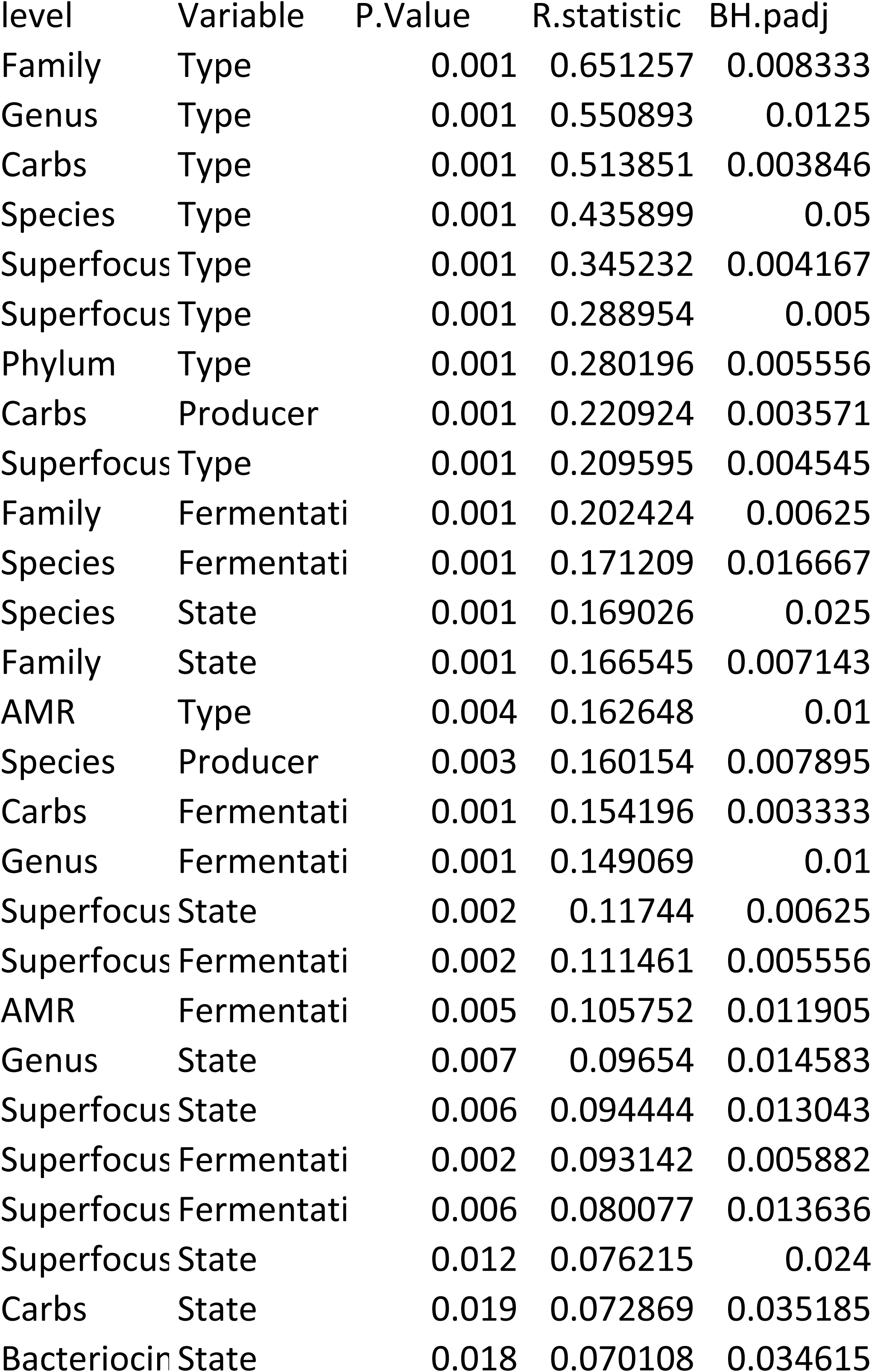
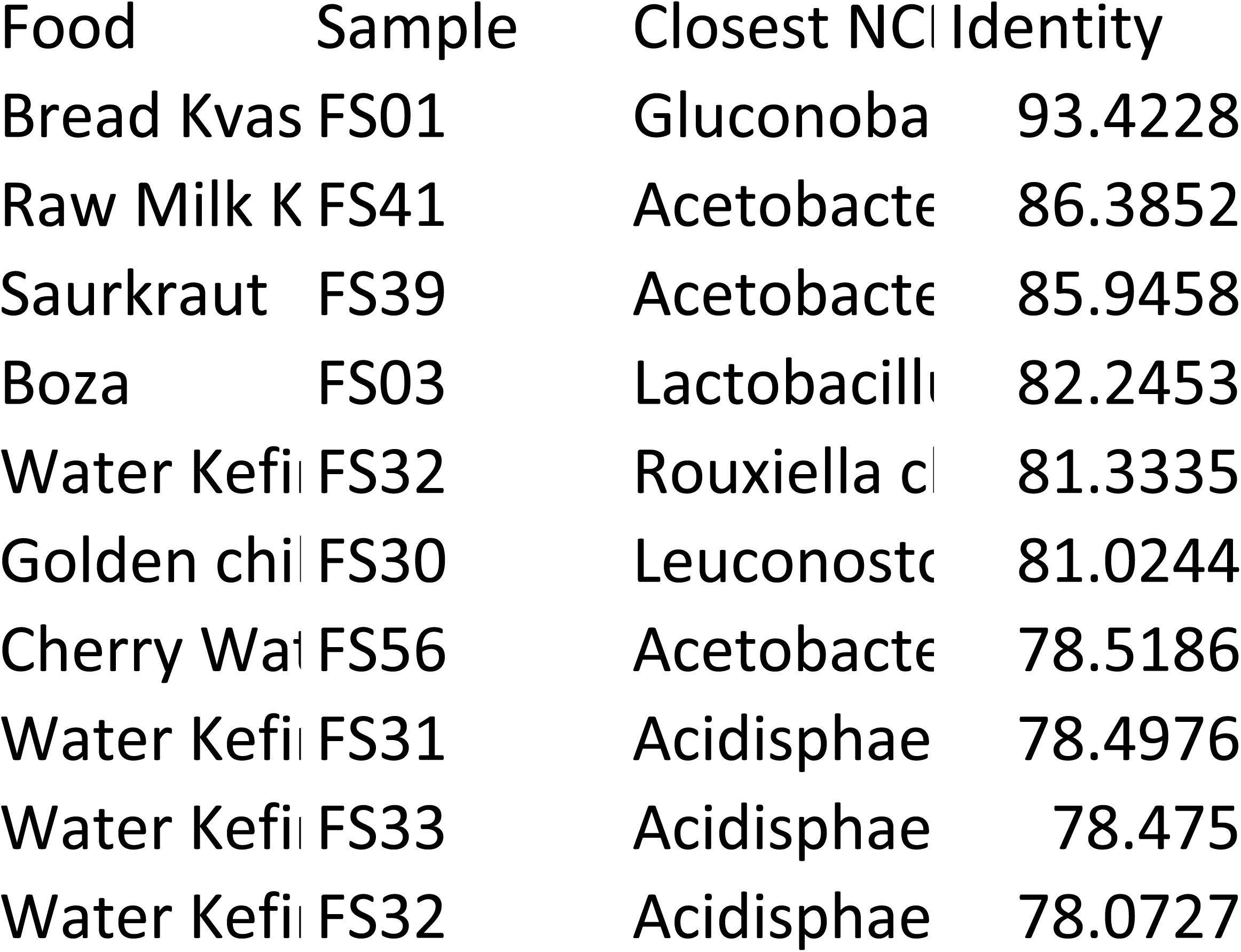

